# Derivation of primary fetal epithelial organoids from cryopreserved human amniotic fluid cells

**DOI:** 10.64898/2026.09.07.749928

**Authors:** Giorgia D’Ariano, Gloria Ji Zhang, Giuseppe Calà, Giuseppe Matteo Carrino, Aeshna Agarwal, Alessandro Mariani, Michela Marinaro, Rodolphe Matias de Sousa, Gianluca Cenedese, Carlotta Camilli, Anna L David, Simon Eaton, Alessandro Filippo Pellegata, Marco Pellegrini, Paolo De Coppi, Mattia Francesco Maria Gerli

## Abstract

Primary fetal human epithelial organoids are canonically derived from tissue samples obtained after termination of pregnancy. Recently, we have demonstrated that these organoids can be consistently generated from amniotic fluid cells isolated prenatally during diagnostic and interventional procedures. Amniotic Fluid-derived Organoids (AFOs) are promising fetal epithelial lung, kidney and small intestine tissue models that bypass some of the ethical and legal constraints associated with obtaining primary fetal tissue. Despite this, the widespread adoption of this technology is limited by the lack of standardised and accessible pipelines for the biobanking, culturing and distribution of organoid-forming amniotic fluid cells. To enable broader AFO use for research and clinical purposes, this work investigates two cryopreservation strategies to optimise AF cell recovery for organoid derivation. In Strategy 1, viable AF cells were sorted before freezing, while in Strategy 2 unsorted AF was frozen, and viable AF cells were sorted for viability after thawing. We present here a comprehensive evaluation of 5 commercially available GMP-compliant freezing media (FM1-5) alongside a standard lab-grade control (FM CT). AFOs were assessed for formation efficiency, morphological characteristics, proliferation capacity, epithelial identity, and tissue type. Our results demonstrate that organoid-forming AF cells can be successfully cryopreserved both pre-and post-sorting. Strategy 1 yielded higher organoid formation efficiency, with AFOs derived from cryopreserved and fresh cells exhibiting comparable expansion potential. Finally, FM4 provided minimal to no decline in survival rate, making it the most effective GMP-grade freezing medium tested. In conclusion, we present two viable cryopreservation strategies adaptable to different laboratory settings and identify optimal GMP-compliant freezing media, to support safe distribution and centralised processing, thereby facilitating scalability and collaborative work on the AFO technology.

**HIGHLIGHTS:**

- Cryopreserved AF cells efficiently generate primary fetal epithelial organoids (AFOs)
- Live cell sorting conducted before cryopreservation retains AFO formation efficiency
- GMP-compliant freezing media show different AF cells cryopreservation ability
- AFOs from cryopreserved AF cells expand well, and retain tissue-specific phenotype

## INTRODUCTION

Organoids are a powerful tool to study human tissue development, homeostasis and disease^1,2^. Human organoids are undergoing extensive clinical testing for *in vitro* disease modelling and as a cell source for regenerative medicine^3^. Primary organoids, established from tissue-resident progenitors, have the ability to recapitulate cellular architecture and physiology of the epithelial tissues, making them valuable platforms for patient-specific modelling and personalised medicine applications. Postnatal primary organoid derivation is mostly conducted from tissue biopsies and discarded surgical resections^4^. Differently, collection of fetal tissue biopsies from continuing pregnancies is considered unfeasible due to high fetal and maternal risks. Therefore, primary fetal organoids have been previously isolated post-mortem from donated fetal specimens collected following termination of pregnancy (TOP). This carries legal and ethical restrictions in several countries and, even when permitted, it may only be accessed in a narrow gestational window depending on the indication (https://cks.nice.org.uk/topics/abortion/background-information/uk-abortion-laws/). Altogether, these implications limit the use of primary fetal organoids in basic science and clinical applications. While a valid alternative platform for studying fetal development is represented by organoids derived from pluripotent stem cells, this has some limitations in personalised prenatal medicin, such as the extensive manipulation and time required for its implementation. In this context, an alternative source of fetal epithelial progenitors is represented by the amniotic fluid (AF)^5^. The AF contains epithelial cells with renal, pulmonary and gastrointestinal identity, released in the fluid during development. Such tissue-specific epithelial cells are able to form primary organoids autologous to the developing fetus, termed amniotic fluid organoids (AFOs)^5^. To do this, fresh AF samples are centrifuged, filtered and sorted for live cell isolation. AF cells are plated in three-dimensional Matrigel droplets and grown in epithelial medium allowing for AFOs formation within 10-14 days. Individual AFO are then picked and expanded clonally, screened for their tissue identity, expanded for multiple passages and routinely cryopreserved without affecting their proliferating potential. AF samples are accessible across the whole second and third trimester of gestation through multiple routine clinical procedures such as amniocentesis and amniodrainage, fetal surgery and during caesarean section birth. Thanks to this accessibility, the AF allows for patient-specific organoid derivation at a broad gestational age range, compatibly with pregnancy continuation. With great translational potential, AFOs could be employed for disease modelling and drug testing, in a timeframe compatible with prenatal intervention. It is therefore crucial to implement strategies to facilitate and support the use of AFO in research and clinical laboratories. A critical step in this direction is the development and optimization of robust AF cell cryopreservation protocols aiming at efficient generation of AFOs post-thawing. Isolation and organoid derivation from fresh AF require access to cell sorting, training in three-dimensional cell culture and is associated with significant costs. Such specialised requirements are often not available in clinical laboratories, in geographically remote areas and in low-and middle-income countries (LMIC). Therefore, optimising samples distribution, biobanking and centralisation of organoid production, might help in addressing these issues^6^. Optimising the methodologies for AF cell banking would provide a readily available source of cells, enabling potential future translation of personalised AF-derived models and cellular products. Nowadays numerous biobanks are dedicated to the storage of perinatal-derived stem/progenitor cells^7,8^. Umbilical cord blood hematopoietic stem/progenitor cells, for example, have been efficiently stored for decades in public and private biobanks for medical research and haematological disease treatments^9,10^. Although AF cell biorepositories represent a valuable resource, AF cryopreservation would also need to be optimized to enable effective AFO formation. This will allow dissemination of the technology and the creation of patient-and disease-specific repositories for future clinical and research use. In this context, primary organoids and tumoroids have been successfully derived from a number of frozen solid human biopsies, including intestine, lung, endometrium, brain and pancreas specimens^11–19^. Organoids from fresh and cryopreserved biopsies have comparable expression of tissue specific markers^14,19^ and responses to hormonal/drug treatments^13,15,16^. Organoid derivation from frozen samples is therefore technically feasible, but this has not yet been reported from the AF. The development and optimization of a robust protocol for cryopreserving AF cells enabling efficient generation of AFOs at thawing is therefore crucial. Overall, AF cell cryo-storage, optimised for effective post-thawing AFO derivation, would enable collaborative research and diagnostic work amongst distant laboratories. In this article we present two freezing strategies for AF cell storage: in strategy 1, live cell sorting was performed before freezing the cells; in strategy 2, the AF cell fraction was collected and frozen as a whole, while live cell sorting was conducted post-thawing. Moreover, we conducted comparative analysis by deriving AFOs from AF cells, cryopreserved in five GMP-compliant commercially available freezing media (FM1-FM5) versus our lab-grade counterpart (FM CT) (Table 2). Overall, here we present an effective strategy for AF cell storage optimised for AFO derivation.

## Materials and Methods

### Freezing media formulations

Six different freezing media (FM) were tested in this work. FM CT consisted of a lab grade medium containing 10% DMSO (Sigma-Aldrich, 472301), 40% FBS (Gibco, 26140079), 50% ADMEM+++ consisting of Advanced DMEM/F12 (Thermo 12634) supplemented with 2mM (1x) Glutamax (Thermo 35050061), 1% Pen/Strep (Thermo 15140122) and 10mM (1x) HEPES (Thermo 15630080). FM1-FM5 represented five commercially available freezing media (Table 1).

### Isolation and cryopreservation of amniotic fluid cells

Human AF samples were collected after informed consent in compliance to all relevant institutional and governmental regulations under ethical approval by the NHS Health Research Authority - London Bloomsbury Research Ethics Committee (REC 14/LO/0863 IRAS ID 133888). All samples were obtained at the University College London Hospital (UCLH) Fetal Medicine Unit (FMU) with informed written patient consent. AF samples were collected during amniodrainage procedures performed as part of standard clinical care for pregnancies with polyhydramnios caused by twin-to-twin transfusion syndrome (TTTS) between 19 and 25 weeks of gestation. This type of samples was selected due to the large volume required to test the multiple conditions included in the study. After collection, fluids were stored at +4°C and processed within 24h. Each sample was split into aliquots and prepared separately following the two strategies outlined below:

*Strategy 1 and Fresh controls:* AF samples were processed as previously described^5^. Briefly, the fluid was passed 70 µm and 40 µm cell strainers and centrifuged at 300 g for 10 min at 4°C. The pellet was resuspended in 50 mL of cold FACS Blocking Buffer (FBB: 1%FBS, 0.5 mM EDTA in PBS) and centrifuged again at 300 g for 10 min at 4°C. Following this, the pellet was resuspended in 2-3 mL of FBB and incubated for 40 min at 37°C with Hoechst 33342 (Sigma-Aldrich, 33342) at a final concentration of 5 µg/mL. Propidium Iodide (PI, Sigma-Aldrich, P4170) was added at final concentration 2 µg/mL and incubated for 5 minutes at room temperature (RT). Viable cells were sorted using a FACS Melody (BD Biosciences) gating for the Hoechst^pos^ / PI^neg^ population. To preserve AF cellular heterogeneity, no gate was applied to forward or side scatter. Cell viability was confirmed through Trypan Blue staining (Sigma-Aldrich, T8154), counting the viable cells in a haemocytometer. Sorted cells were then divided into seven aliquots, corresponding to our seven experimental groups. One aliquot was used as Fresh control and plated immediately in organoid culture conditions as detailed below. The other six aliquots were frozen using six different freezing media. In the case of FM CT, cells were resuspended in 1:1 ice-cold ADMEM+++ and freezing solution (20%FBS, 80% DMSO). For FM1-FM5, 1 mL of cold medium was directly added to the cell pellet dropwise. Cells resuspended in freezing medium were then transferred to cryovials and moved into a slow-cooling box (Mr Frosty, Thermo Scientific™ 5100-0001), kept at −80°C overnight and transferred to LN_2_ the following day.

*Strategy 2:* Unsorted AF samples were taken through a series of washes to remove cellular and acellular debris. Briefly, AF was strained over 70 µm and 40 µm cell strainers and centrifuged at 300 g for 10 min at 4°C. The cell pellet was then resuspended in 60 mL ice-cold FBB and divided in six aliquots. Each aliquot was centrifuged at 300 g for 10 min at 4°C. This last step was repeated, for a total of three washes. After the third centrifugation step, supernatant was removed, and the cell aliquots were frozen in the six different freezing media as described for Strategy 1 (Table 1).

### AF cells thawing

Cells frozen with Strategy 1 and Strategy 2 were thawed on the same day, after 7-14 days of cryopreservation in LN_2_. For both strategies the cryovials were equilibrated on dry ice and then placed at 37 °C allowing partial thawing. 1mL of ice-cold ADMEM+++ was added dropwise to the vials to avoid osmotic shock. After a gentle resuspension, the vial contents were rapidly transferred to 15 mL Falcon tubes and 8 mL of ice-cold ADMEM+++ were added. Samples were then centrifuged at 300 g for 5 min at 4 °C. To eliminate as much freezing medium as possible, a second washing was performed. For *Strategy 1*, thawed cells were plated and cultured for organoid derivation as in the Fresh controls (see “AFO derivation and expansion”). For *Strategy 2*, the pellet was resuspended in 2-3 mL of FBB, incubated for 40 min at 37°C in Hoechst 33342 (5 µg / mL) and 5 min at RT in PI (2 µg / mL). Live cells were sorted using a FACS Melody (BD Biosciences) gating for Hoechst^pos^ / PI^neg^. Viability was confirmed through Trypan Blue staining and manual counting. Sorted cells were centrifuged at 300 g for 5 min at 4 °C and plated as described for Strategy 1 and Fresh control as described below.

### AFO derivation and expansion

AFO were derived and expanded as previously described^5^. Briefly, cells were centrifuged at 300 g for 5 min at 4 °C, resuspended in 30µL of cold Matrigel (Corning, 354230) and plated as pellets, onto a pre-warmed 24-well plate. The plate was then incubated for 20 min at 37° to allow Matrigel gelation. The cells were then cultured in a chemically defined expansion medium (AFO expansion medium, Table 2) supplemented with 10μM Rho-kinase inhibitor (ROCKi; Tocris, 1245) and 2μl/mL Primocin (Invivogen, NC9141851) for the first 2 days of culture. Medium was replaced every 3-4 days.

14-20 days after the initial plating, individual organoids were manually picked under the microscope and clonally expanded. Each picked organoid was transferred to a 0.5 mL tube pre-coated with 1% BSA, resuspended in TrypLE and incubated for 3-7 minutes at 37°C. Following digestion, 400 μL of ice-cold Advanced ADMEM+++ was added to dilute the TripLE. Finally, the organoids were mechanically dissociated by pipetting and the suspension was centrifuged at 300g for 5 minutes at 4°C. After centrifugation, the pellet was resuspended in 20 μL of cold Matrigel and then plated into a pre-warmed 48-well plate. The plate was incubated at 37°C for 20 minutes, then AFO expansion medium (Table 2) supplemented with 10μM ROCK inhibitor. The medium was changed every 3 days without adding ROCKi. After 10 to 14 days, depending on their number and size, the organoids were split into either a 24-well or 12-well plate. For passaging, medium was replaced with ice-cold ADMEM+++. The Matrigel droplets were then broken down and collected into a 15-mL tube on ice. The organoids were washed with 10 mL of cold ADMEM+++ and centrifuged at 300g for 5 minutes at 4°C. After centrifugation, the supernatant was carefully aspirated and 300 μL of TrypLE were added to each tube to facilitate cell dissociation. The tubes were incubated at 37°C for 3-5 minutes. After enzymatic digestion, organoids were resuspended in 1 mL of ADMEM+++ and mechanically disaggregated into single cells by pipetting. Next, cold ADMEM+++ was added to a total volume of 10 mL to further dilute TryplE and the samples were centrifuged at 300g for 5 minutes at 4°C. The cell pellet was then resuspended in Matrigel and plated. The plate was incubated for 20 minutes at 37°C to allow the Matrigel to solidify. Finally, AFO expansion medium (Table 2), along with 10μM ROCK inhibitor was added and the medium was changed every 3-4 days.

### Organoid formation efficiency and morphometric analyses

Organoid formation efficiency was determined at day 14 after AF cells plating by counting the number of organoids at P0.

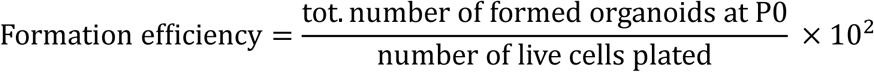

Organoid area and circularity were determined for each organoid formed at P0 by measuring the perimeter in 5x or 10x magnification transmitted light microscope images (Zeiss Axio Observer A1) using the ImageJ software.

### Whole mount immunofluorescence

Culture medium was replaced with 1 mL of Cell Recovery Solution. The plates were then placed on ice for 45 minutes to allow Matrigel to melt. The solution containing the organoids was then transferred into Eppendorf tubes pre-coated with 1% BSA and centrifuged at 200 RCF for 2 minutes. The supernatant was aspirated carefully and 1 mL of 4% PFA was added at room temperature (RT) for 30 minutes. After the incubation, organoids were spun for 1 minute, washed 3 times with PBS for 5 minutes. For whole-mount immunostaining the organoids were subjected to blocking/permeabilization with a solution of PBS-Triton X-100 0.5% with 1% BSA for 1 hour at RT. The samples were then incubated in the blocking/permeabilization solution containing the primary antibodies for 24 hours at 4°C in agitation. After the incubation with primary antibodies, organoids were washed with PBS-Triton X-100 0.2%. Next, the organoids were incubated with secondary antibodies and Hoechst in the blocking/permeabilization solution overnight at 4°C, in agitation. After secondary antibody incubation, the organoids were washed again and resuspended in PBS for confocal imaging. The list of primary and secondary antibodies used is available in Table 3. Immunofluorescence images of the whole-mount staining were acquired using a Zeiss LSM 710 confocal and Zeiss LSM 980 confocal microscopes with x10 and x20 immersion and air objectives. ImageJ software was used to generate z-stack projections and perform image analysis.

### Statistical analysis

Statistical analysis was conducted on at least three biological replicates. Statistical significance was measured with GraphPad Prism v.10.4.1 software using paired t-test when comparing two groups and one-way ANOVA when comparing more than two groups (Fresh vs. thawed groups), as reported in the figure legends.

## RESULTS

### Viable AF cells recover from cryopreservation with or without live cell sorting

Fresh AF samples were collected from 4 patients (19-25 gestational age weeks) undergoing amniodrainage for twin-to-twin transfusion syndrome (Supplementary Table 1). Each AF sample was processed for both fresh plating and cryopreservation utilising two different strategies (Figure 1). Live cells were isolated via FACS and either freshly plated as control or immediately frozen for Strategy 1. In Strategy 2, the entire AF cell fraction was frozen within 24 hours from collection, and live cell sorting was performed right after thawing. For each strategy we tested six different freezing media formulations as listed in Table 1. In particular, we tested a standard lab-grade freezing medium commonly used for cryopreservation of organoids (FM CT), against five GMP-compliant, commercially available freezing media marketed for stem cells cryopreservation (FM1 to FM5). FM1, FM3, and FM4 contain dimethyl sulfoxide (DMSO) as cryoprotecting agent, while FM2 and FM5 are labelled as DMSO-free. Therefore, we evaluated thirteen experimental conditions: i) fresh plating after live cell sorting; ii) plating after freezing using Strategy 1 with six different freezing media (six conditions); iii) plating after freezing using Strategy 2 with the same six freezing media (six conditions). Accordingly, the initial AF volume was divided into thirteen equal fractions to accommodate each condition. For Strategy 1, an AF volume consisting of seven fractions was sorted for Hoechst^pos^ / PI^neg^ staining to isolate viable cells (Figure 2A and 2B) which were further partitioned: 4-5 x 10^4^ Hoechst^pos^ / PI^neg^ cells were plated fresh as control for organoid derivation or frozen in each of the six FM. For Strategy 2, the remaining AF volume was passed through 70 µm and 40 µm cell strainers in order to remove clumps and large debris, resuspended in FBB, further divided into six aliquots, and each was washed three times in FBB. Each pellet was then frozen in one of the six FM. Fresh-plated viable AF cells were used as control for both Strategy 1 and Strategy 2. Cells were thawed after 7-14 days, embedded in Matrigel and cultured in AFO Expansion Medium for organoid derivation (Table 2). For both strategies, thawed cells were firstly washed twice in ADMEM+++ to remove the freezing medium and plated right away in the case of Strategy 1. For Strategy 2, thawed cells were stained and sorted for Hoechst^pos^ / PI^neg^ (Figure 2C) and then plated. To assess which conditions had the most efficient cryopreservation properties, we compared the percentage of live cells recovered after thawing (Figure 2D). As expected, a reduction in the number of live cells was observed for all FM and for both strategies. Utilising Strategy 1, the average percentage of live cells retrieved at thawing varied between 44.19% (FM1) and 27.29% (FM3). With Strategy 2, instead, we observed a slightly lower range of recovered live cells: between 26.45% (FM CT) and 20.06% (FM5). No statistically significant difference was observed when comparing the six FM or the two strategies.

**Figure 1.**
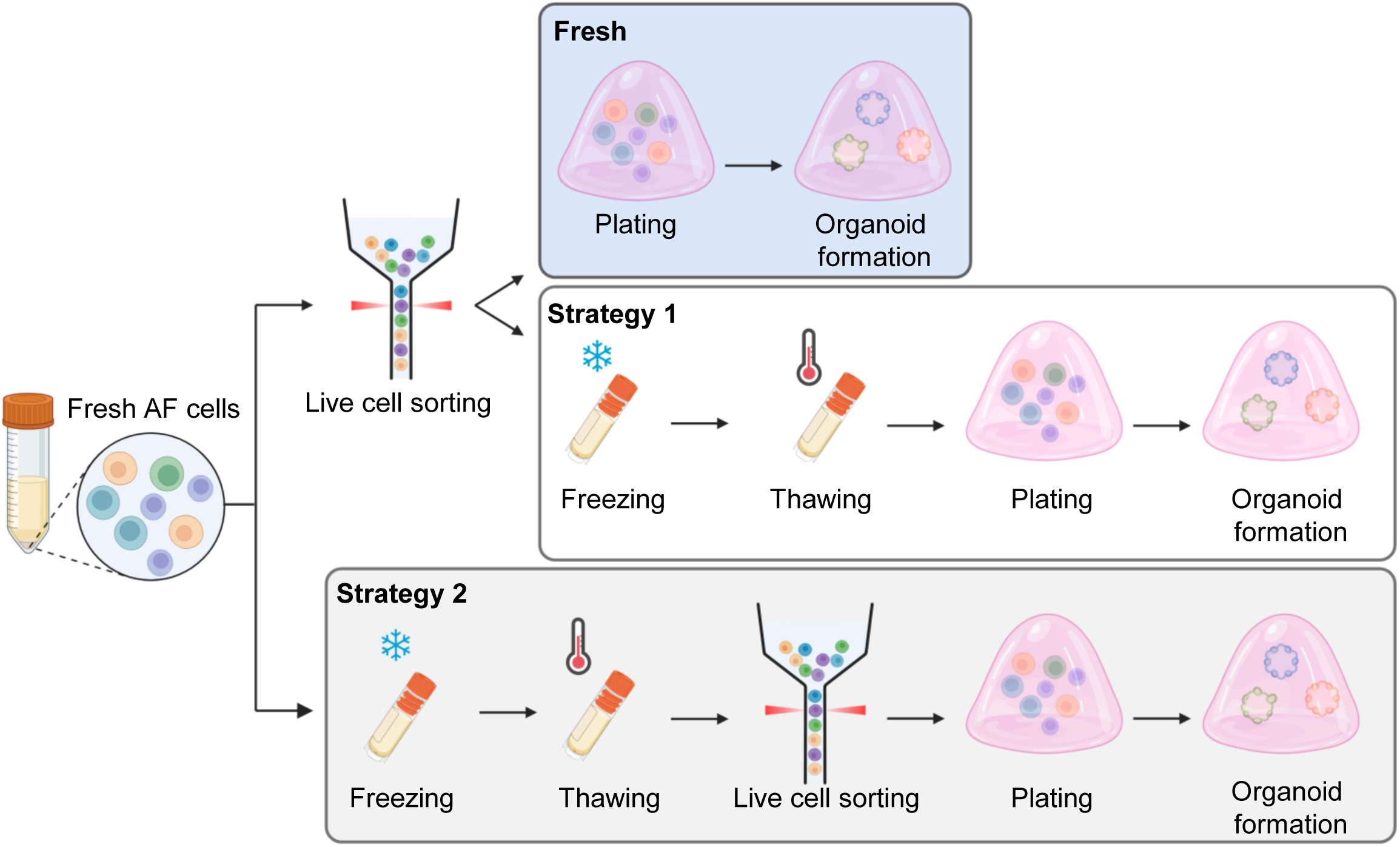
AF cells freezing strategies. Graphical abstract of experimental design. Images created with BioRender.

**Figure 2.**
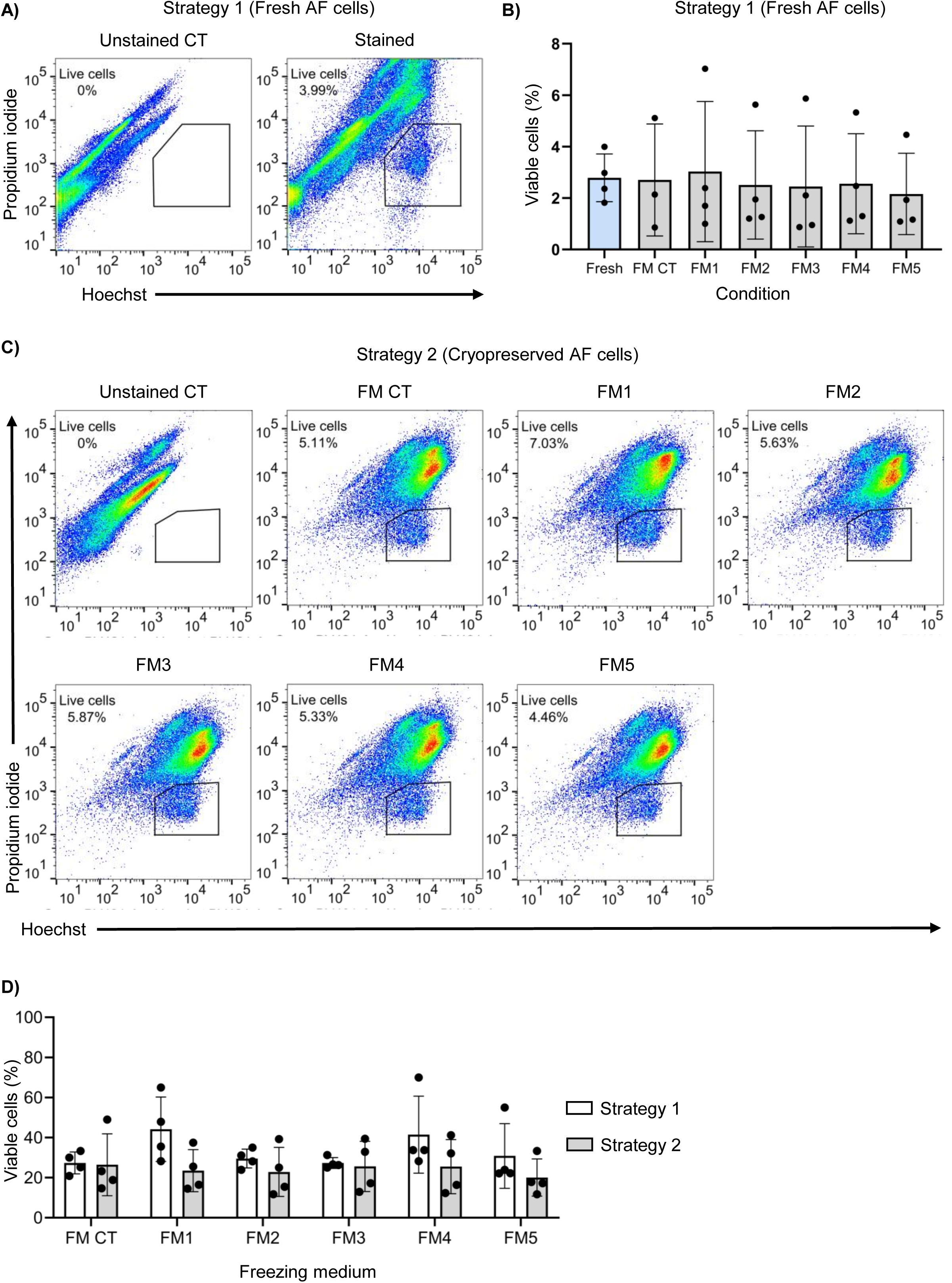
AF cells can be cryopreserved pre-and post-live cell sorting. (A) Representative FACS plot showing the gating strategy utilised to isolate the live cell fraction from fresh AF samples (negative for Propidium Iodide and positive for Hoechst); (B) Percentage of live cells sorted from n = 4 independent fresh AF samples (n=3 for FM-CT). Cells were counted with Trypan blue; mean ± s.e.m., Fresh and each FM were compared with one-way ANOVA with multiple comparisons (non-significant); (C) Representative FACS gating utilised for live cell selection post-thawing (Strategy 2); (D) Percentage of viable cells after thawing, relative to the total number of cells frozen. Cells were counted with Trypan blue after thawing for Strategy 1 and after live cell sorting for Strategy 2; mean ± s.e.m., Strategy 1 and Strategy 2 were compared with multiple paired t-test (non-significant).

### Cryopreserved AF cells form AFOs

To determine whether cryopreserved AF cells maintain their ability to generate AFOs, we compared the organoid formation efficiency between the different conditions (Figure 3A). All freshly plated samples led to organoid formation with mean efficiency of 0.04%. Samples frozen utilising Strategy 1 displayed non statistically significant difference in organoid derivation efficiency, although in the case of FM2 and FM5 1 out of 4 samples (25%) showed no organoid formation. In Strategy 2 we observed an overall drop in the percentage of samples forming organoids when compared to the fresh control. Indeed, AFO formation was observed in 3 out of 4 samples when using FM CT, FM3 and FM4, 2 out of 4 samples when using FM5 and 1 out of 4 samples when FM1 and FM2 were utilized. These results suggest that AFOs can be derived from AF cells cryopreserved with different freezing media. Furthermore, when AF cells were cryopreserved employing Strategy 1, AFO formation efficiency was not affected regardless the FM that was used, with the exception of FM2 and FM5. AF cells cryopreserved utilising Strategy 2 were still able to form AFOs but with a lower efficiency. Because of these observations and of the limited number of AFO-forming samples when using Strategy 2, we decided to focus and further investigate Strategy 1 to efficiently freeze and store AF cells. AFOs usually start forming between day 7-10 post AF cell seeding and are picked within 2-3 weeks when reaching a diameter size of at least 100µm (^20^). To assess whether AFOs derived from cryopreserved AF cells maintained a comparable formation growth potential at isolation (P0), we measured organoid area at days 7 and 14 (Figure 3B and 3C). In both thawed and fresh conditions, AFOs could already be observed from day 7 with comparable size and consistently grew in the following week. This indicates that AFOs derived from thawed AF cells form and grow in a timeframe comparable with the fresh control, and similarly to our previous findings^5^.

**Figure 3:**
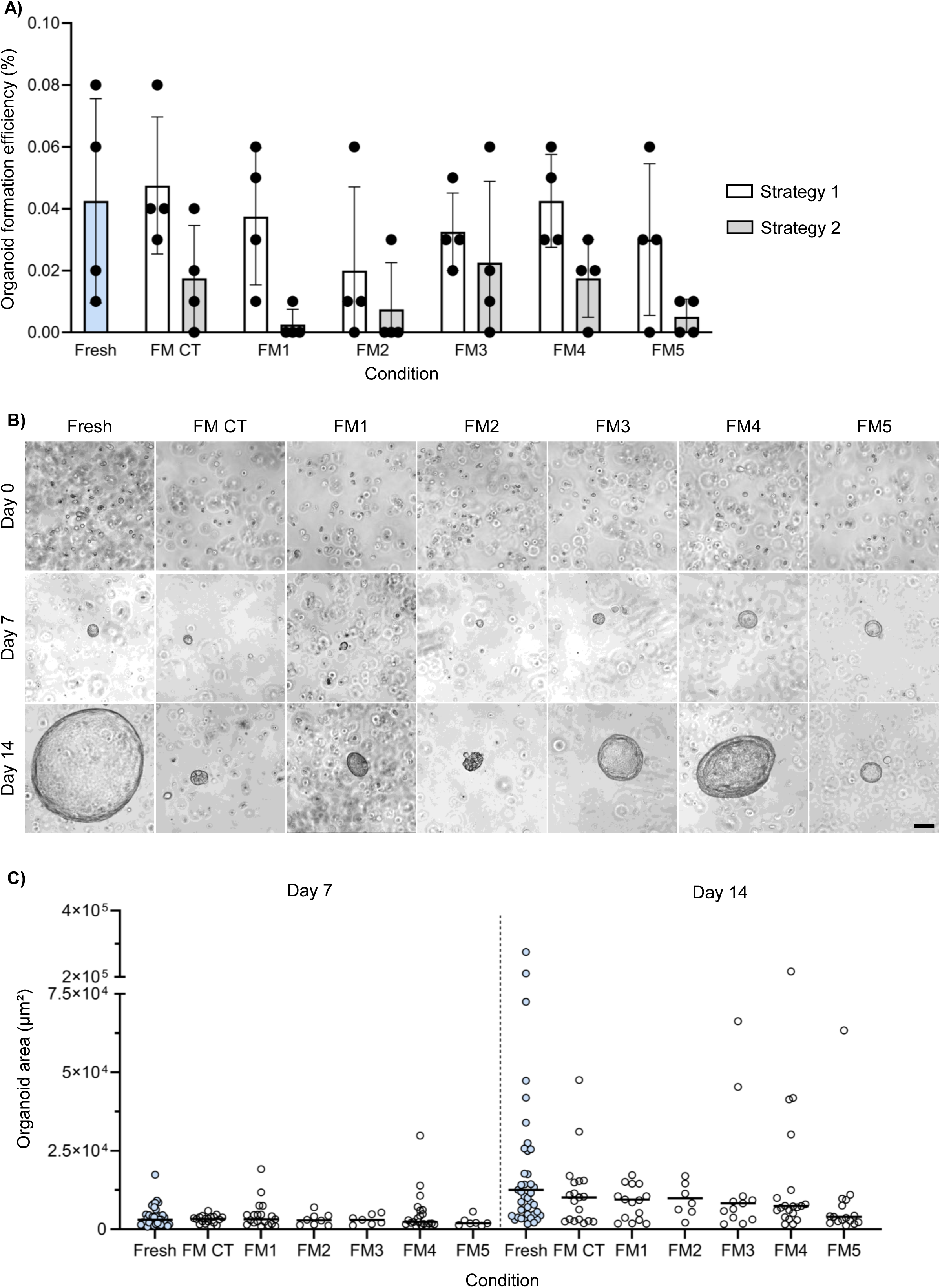
AFOs derivation from cryopreserved AF cells. (A) Organoid formation efficiency at day 14 post seeding of fresh and cryopreserved AF cells. (n = 4 independent biological samples). mean ± s.e.m., For fresh vs cryopreserved comparison one way ANOVA was used (non-significant). For Strategy 1 vs. Strategy 2 comparison, multiple paired t test was used (non-significant); (B) Brightfield images of AFOs from fresh and Strategy 1 conditions. Images were acquired at passage 0 on day 0, 7 and 14 post cells seeding (scale bar = 100µm); (C) Quantification of organoid area at passage 0 of fresh and Strategy 1 conditions at day 7 and day 14 post seeding (each dot represents 1 organoid from n=4 independent biological samples, mean is shown). Fresh condition was compared to each FM using one-way ANOVA (non-significant).

### Organoids from cryopreserved and fresh AF have comparable phenotype

AFOs have an epithelial phenotype and can exhibit multiple morphologies (cystic, compact, budding) at isolation^5^. To confirm that cryopreservation does not affect the intrinsic heterogeneity of AF cells we assessed the AFOs morphology at day 14 post-plating. All tested conditions displayed formation of cystic and compact organoids (Figure 4A). Consistent with our previous observations, budding organoids were not detected, as this morphology only presents between GA 16-18 (^20^), whereas the samples in this study ranged 19-25 GA. Freshly cultured AF cells formed 24% of cystic organoids and 76% of compact ones (Figure 4B). Comparable percentages were observed for FM1 (23.53% cystic, 76.47% compact), FM2 (25% cystic, 75% compact), FM3 (25% cystic, 75% compact), FM4 (24% cystic, 76% compact). A slightly higher proportion of compact organoids was instead observed in FM CT (89.46%) and FM5 (82.35%). As an additional morphometric parameter, we measured the distribution of organoid circularity (Figure 4C). We detected no significant differences between AFOs derived from fresh or cryopreserved AF cells, suggesting that morphological properties are maintained between experimental conditions. Furthermore, whole-mount immunofluorescence on clonally expanded lines confirmed that AFOs derived from thawed AF cells maintained an epithelial phenotype as indicated by the presence of the epithelial adhesion proteins EPCAM (Figure 4D) and ECAD (Figure 4E).

**Figure 4:**
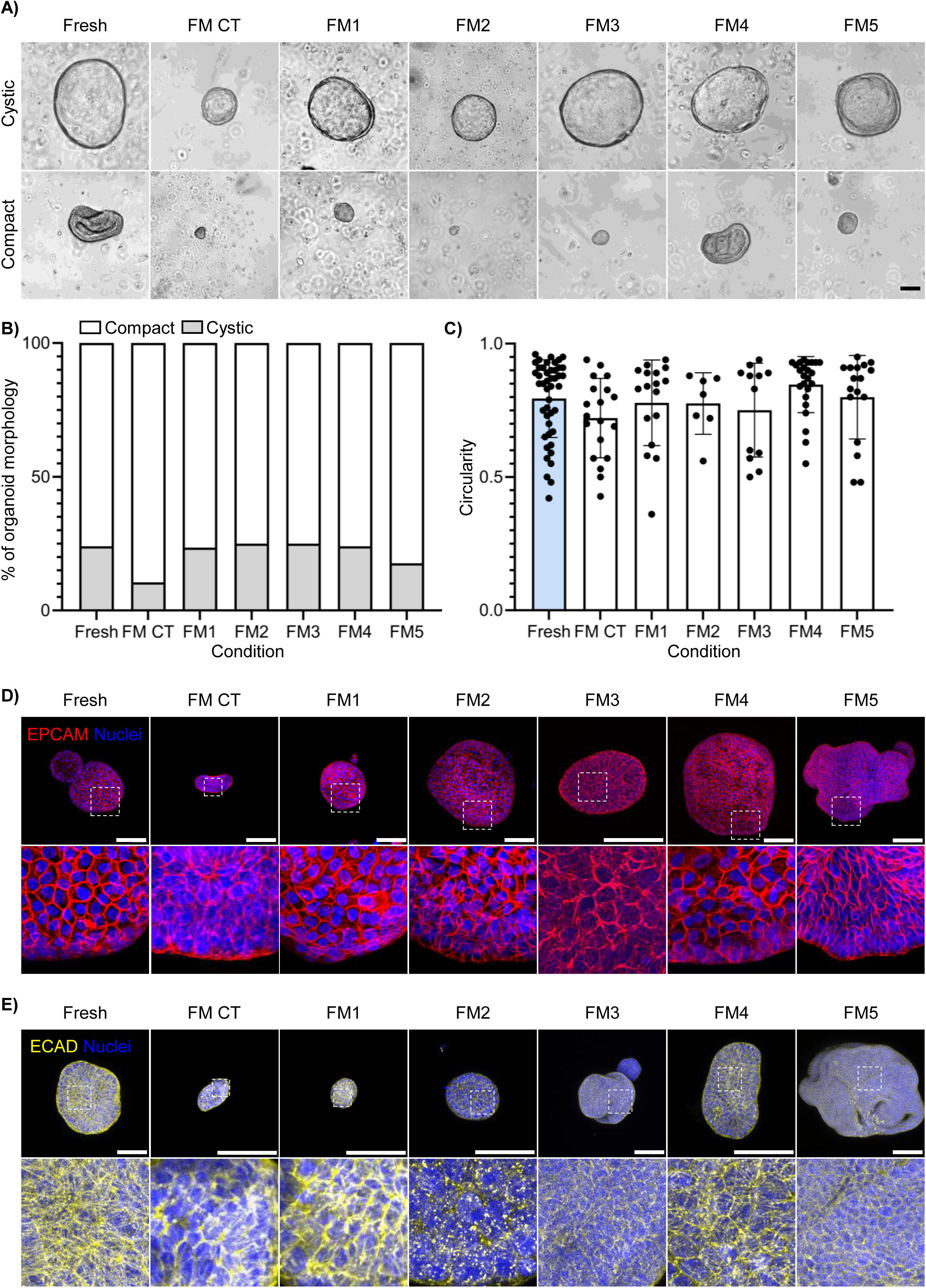
AFOs from AF cells cryopreserved through Strategy 1 have a phenotype comparable to fresh controls. (A) Brightfield images of cystic and compact AFOs at passage 0, 14 days post seeding, from fresh and cryopreserved samples (scale bar = 100 µm); (B) Quantification of cystic and compact AFOs derived from fresh and cryopreserved AF cells. Compact and cystic organoids were counted for each condition at day 14 post-seeding (passage 0); percentage is relative to the total number of formed organoids (n = 4 independent biological samples). Fresh vs FM comparison was performed with one-way ANOVA (non-significant); (C) Circularity measurement of AFOs derived from fresh and cryopreserved conditions at day 14 post-seeding (passage 0). Each dot represents 1 organoid from n = 4 independent biological samples. mean ± s.e.m., Fresh vs FM comparison was performed with one-way ANOVA (non-significant). (D) and (E) whole mount immunofluorescence staining of epithelial markers EPCAM (D) and ECAD (E) counterstained with Hoechst in AFOs derived from fresh and cryopreserved samples (scale bar = 100 µm).

### Organoids derived from amniotic fluids cryopreserved with GMP-grade freezing media can be expanded for multiple passages

After AFOs form and grow at passage 0, single organoids are manually picked and enzymatically dissociated for clonal expansion. We observed that picking AFOs that are at least 100 µm in their longest diameter show optimal recovery post-dissociation (^20^). Once AFOs recover from picking, they can be expanded for multiple passages via mechanical or enzymatic dissociation. However, different AFO clonal lines derived from the same fluid do not have the same proliferative potential, therefore a proportion of the organoids that form at passage 0 fails to expand. Consequently, in addition to checking AFO derivation efficiency, it is fundamental to determine if cryopreservation affects the subsequent proliferative potential of the organoids. To address this, we picked and clonally expanded AFOs derived from fresh and thawed samples and then measured the proportion of clonal lines that could be efficiently expanded for multiple passages. First, we picked all organoids resulting to have diameter equal or longer than 100 µm at 14 days post seeding (Supplementary Figure 1A). The average percentage of organoids over 100 µm per sample varied across the different conditions, spanning between 12.82% (FM1) and 62.22% (Fresh) (Supplementary Figure 1B). We then proceeded to expand the selected AFOs, therefore establishing the clonal lines (Figure 5A). AFO lines survival for each sample was assessed across three different time points: passage 2 (P2), passage 4 (P4) and passage 6 (P6) (Figure 5B, Supplementary Figure1C and 1D). By P2, an expected drop in survival was observed in all groups, with FM2 showing the most pronounced decrease, falling below 20%. FM CT, FM1 and FM3 maintained intermediate survival (∼30-40%), while Fresh control, FM4, and FM5 retained relatively higher viability (∼50-70%). As passaging progressed to P4, minimal to no decline in survival was observed for FM4 and Fresh group, while FM1, FM2, FM3 and FM5 stabilised between 15-35% of survival. Notably, the FM CT group exhibited complete loss of survival. By P6, survival percentages remained consistent with those at P4, suggesting a plateau in viability loss after initial declines. The higher robustness of the FM4 group is also indicated by the fact that it is the only formulation allowing AFO expansion until at least P6 of 3 out of 3 AF samples in the study, consistently to what was observed with the Fresh controls (Figure 5C). We also compared the cell proliferation rate of the AFO lines amongst the different groups (Supplementary Figure 1E, 1F and 1G). Comparable levels of Ki67⁺ cells were observed at P2 with no statistically significant variations. At later passages (P4 and P6), the percentages of Ki67⁺ cells remained stable across the experimental groups. These findings indicate that AFOs derived from AF cells frozen with Strategy 1 can be successfully expanded for multiple passages.

**Figure 5:**
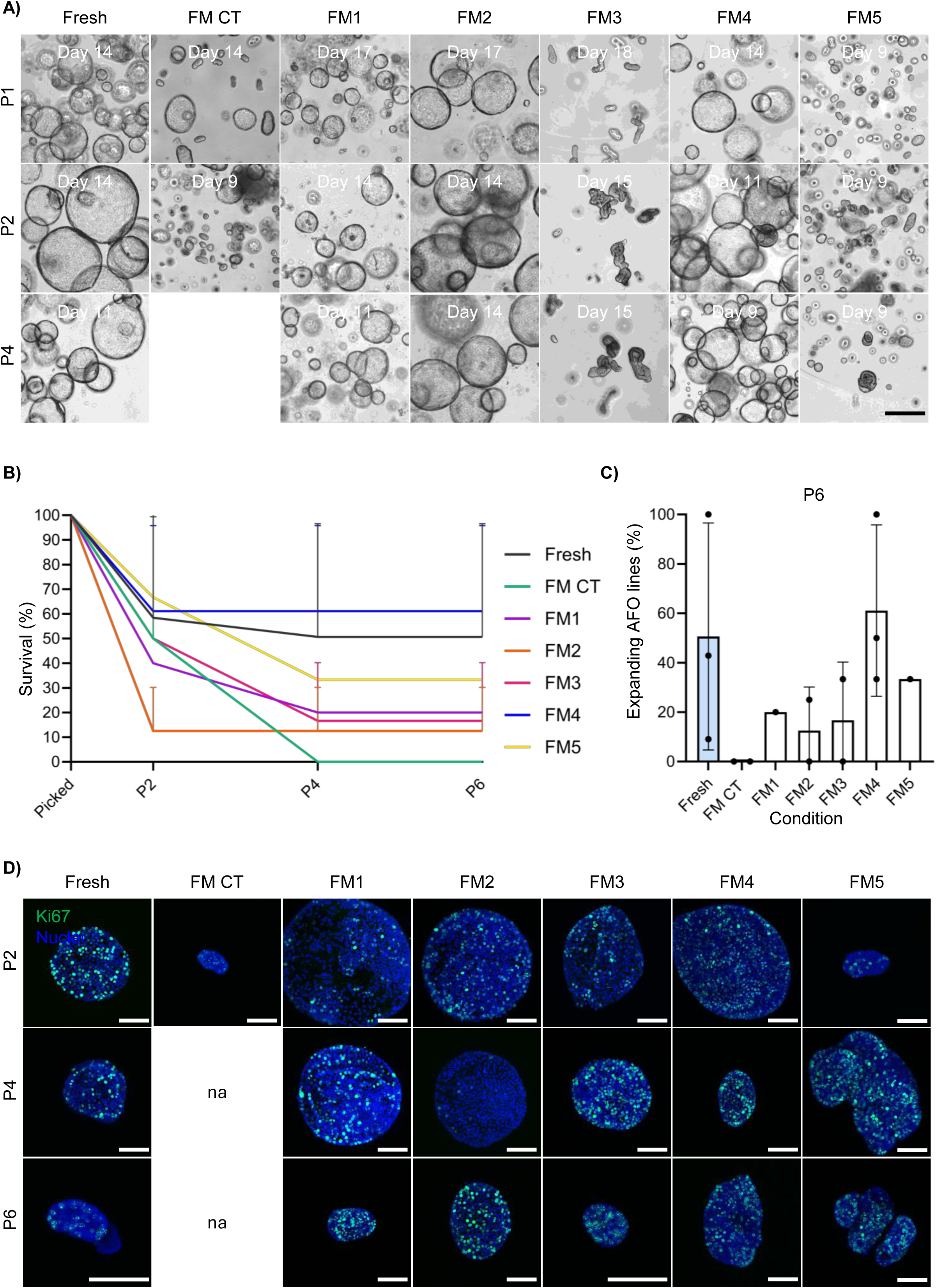
AFOs from AF cells cryopreserved through Strategy 1 can be expanded for multiple passages. (A) Brightfield images of expanding AFOs from fresh and thawed cells at passage 1, passage 2 and passage 4 (scale bar = 100 µm). (D) and (E) percentage of AFO lines that could be expanded up to passage 2 and passage 4, relative to the number of picked organoids. mean ± s.e.m.; (B) Survival curve showing the percentage of AFO lines that could be expanded for each passage (n = 3 independent biological samples). s.e.m. is shown; (C) percentage of AFO lines that could be expanded up to passage 6 (corresponding to P6 in figure 5B). mean ± s.e.m. (D) whole mount immunofluorescence staining of proliferation marker Ki67 counterstained with Hoechst (scale bar = 100 µm).

In comparison, Strategy 2 also resulted to be less efficient in terms of organoid formation efficiency (Figure 3A). At day 14 post seeding we identified pickable organoids in all conditions, with the only exception for the FM5 group (Supplementary Figure 2A). We proceeded to attempt clonal expansion of such AFOs. Consistently with what observed for Strategy 1, none of the lines derived from the groups FM CT and FM2 could be expanded beyond P1 and P2 respectively (data not shown). AFOs of the groups FM1, FM3 and FM4 expanded for multiple passages (Supplementary Figure 3B). AFOs from FM1, FM3 and FM4 displayed at P6 a percentage of Ki67^+^ proliferating cells consistent with those observed in the fresh control and Strategy 1 samples (Supplementary Figure 2C and 2D and Figure 5F). This indicates that while Strategy 2 may decrease the number of derivable AFOs, its use does not affect their proliferative potential.

### AFO from cryopreserved AF cells manifest kidney and lung identities

AFO manifest distinct tissue identities: kidney (KAFO), lung (LAFO), small intestine (SiAFO). While both LAFOs and KAFOs can be derived from most AF samples, we observed the presence of SiAFOs in a narrow GA window (16–18 GA weeks) (^20^). Here, to exclude that cryopreservation affects the intrinsic AF cells heterogeneity and consequently the one of AFOs’, we applied a tissue identification workflow to all the AFO lines that could be expanded at least until P4. We utilised an immunofluorescent staining panel based on tissue-specific markers, to investigate the tissue identity of the AFO derived from cryopreserved AF cells. Briefly, CDX2^pos^ AFO are considered small intestine (SiAFO); CDX2^neg^ / PAX8^pos^ /NKX2-1^neg^ are considered kidney AFO (KAFO), whereas CDX2^neg^/PAX8^neg^ / NKX2-1^pos^ are lung AFO (LAFO). Our staining panel highlighted presence of LAFO and KAFO lines in the freshly plated controls (Figure 6A and 6B). LAFOs and KAFOs were also observed in the FM4 and FM5 groups while in the other conditions either KAFOs (FM1, FM2) or LAFOs (FM3) were not observed. As expected due to the GA of our samples, no SiAFOs were observed in any condition.

**Figure 6:**
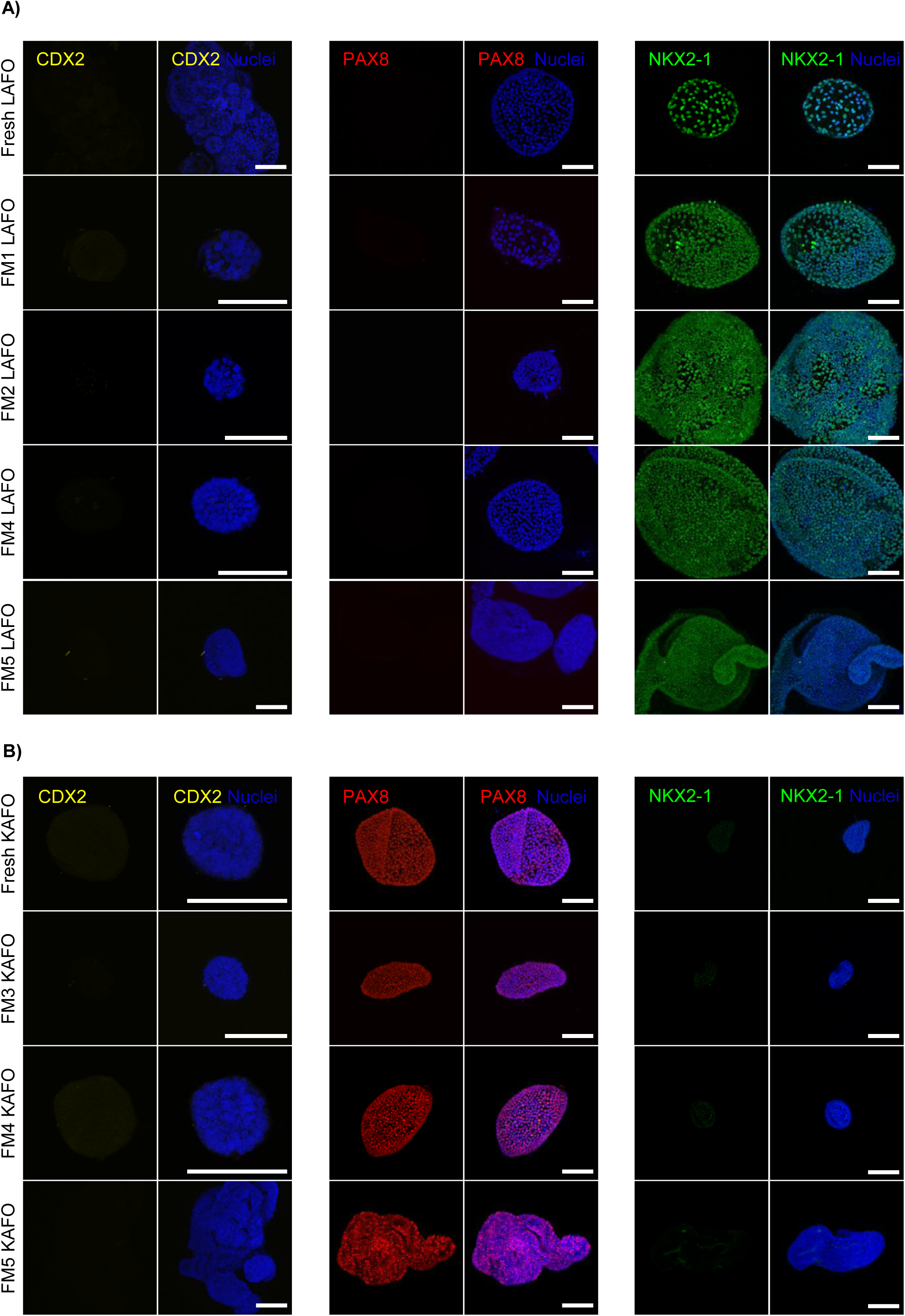
AFOs from AF cells cryopreserved through Strategy 1 can have kidney or lung identity. Whole mount immunofluorescence staining of CDX2, PAX8 and NKX2-1 counterstained with Hoechst on LAFO (A) and KAFO (B) (scale bar = 100 µm).

## DISCUSSION

This study demonstrates that AF cells can be efficiently cryopreserved while retaining their ability to generate expandable fetal epithelial organoids. In this work we evaluated two cryopreservation strategies and compared five GMP-compliant freezing media against our previously reported lab-grade freezing medium (FM CT). In Strategy 1, AF cells were sorted for viability and then cryopreserved, whereas in Strategy 2, unsorted AF cells were cryopreserved and subsequently sorted for viability following thawing. While both cryopreservation strategies allowed recovery of viable AF cells, each freezing media yielded distinct outcomes for cell viability and downstream potential of AFO expansion. Our data indicate that freezing AF cells after live cell sorting (Strategy 1), rather than before (Strategy 2), improved cell recovery and AFO formation efficiency. FM CT is routinely used for cryopreserving established organoid lines while preserving their proliferative potential^5^. However, here we found that FM CT is inefficient in maintaining the ability of cryopreserved AF cells to expand AFOs over P2. Our results indicate that FM4 outperformed all the other formulations in supporting stable AFO expansion over multiple passages, suggesting it as preferred GMP-compliant medium for future biobanking applications. In contrast, FM1, FM2, FM3 and FM5 showed a lower but consistent survival rate. Moreover, both LAFO and KAFO lines were successfully expanded from cryopreserved AF cells, except in FM1, FM2 and FM3. Although this suggests that these media may differentially impact the viability or the expansion capacity of specific progenitor populations, we cannot exclude an observational bias due to the small sample size. In this work, SiAFOs where not observed either in cryopreserved AF cells or fresh samples. This is consistent with our previous observations (^20^), showing SiAFO occurrence only from samples at 16 to 18 GA weeks, while our samples span from 19 to 25 GA weeks. Although Strategy 1 showed superior outcomes, this relies on the use of a cell sorter that might not be always accessible. On the other hand, freezing unsorted AF cells (Strategy 2) resulted in lower AFO formation efficiency and a reduced proportion of samples capable of producing expandable organoid lines. We hypothesize that the elevated concentration of dead cells, cellular debris and urea crystals, accounting for 95-99% of the identities present in the unsorted AF^5^, might interfere with the cryoprotectant agent loading into cells, therefore limiting its effect. This might also explain the higher viability and organoid formation efficiency observed adopting Strategy 1. While both strategies expose cells to the combined stress of FACS and cryopreservation/thawing, the sequence in which these are applied is likely to influence the overall cellular response. We cannot exclude that reduced performances observed with Strategy 2 come from the cumulative stress of thawing followed by FACS-related shear stress, which might be more detrimental than the reverse order used in Strategy 1. Nonetheless, we found that AFO established from Strategy 2 retained their proliferative potential, suggesting that this approach may still be valid in settings without immediate access to a cell sorting facility. The ability to store AF cells for later processing offers a logistical advantage, enabling centralised AFO generation and biobanking. This is particularly relevant for those centres located in remote or in LMICs, where the infrastructure or expertise required for organoid derivation might not be available. Cryopreservation enables sample collection to be physically and temporally decoupled from downstream processing, facilitating collaborative work, centralised biobanking and remote drug screening. This is particularly relevant for rare fetal diseases, where patient cohorts are small and sample availability is usually limited ^21,22^. AF cell biobanks optimised for AFO derivation could serve as autologous sources of epithelial stem/progenitor cells for future therapeutic applications. While we tested five GMP-compliant freezing media, we acknowledge that our pipeline is still far from being clinical grade. Further efforts will be required, in the future, to progress towards the sole use of GMP reagents, to enable the translation of the AFO technology. In conclusion, this work provides a robust pipeline for the cryopreservation of AFO-forming AF cells. We demonstrated the feasibility of both pre-and post-sorting freezing and identified an optimal GMP-grade freezing medium. Overall, the cryopreservation methods outlined in this article facilitate AF samples sharing and improve accessibility and reproducibility of the AFO technology, advancing its applications in prenatal medicine and fetal disease modelling

### Limitations of the study

This study focused on LAFO and KAFOs, as SiAFOs are restricted to a narrow gestational window not extensively represented in our current cohort. Additionally, while we demonstrated phenotypical equivalence with AFO derived from fresh AF and fetal tissue-derived organoids. Future work should explore whether this is maintained after AFO derivation cryopreserved AF, and upon extended culture. Finally, this study only presents the test of GMP-grade freezing media, while future work will be required to bring the entirety of the AFO-generation and culture pipeline to GMP standards.

## Acknowledgments

MFMG laboratory is supported by the Academy of Medical Science (Springboard Award), Medical Research Council (New Investigator Research Grant UKRI3681) the UCL TRO (Therapeutic Acceleration Support Call 10), CDH UK and the Rosetrees Trust. This work was made possible thanks to an EMBO Scientific Exchange Grant (11127) awarded to GDA. GDA is recipient of the Tor Vergata PhD program in Tissue Engineering and Remodeling Biotechnologies for Body Functions. GJZ is recipient of the UCL-China Scholarship Council Joint Research Scholarship. RMdS is supported by a CDH UK research grant and by a “Prix de l’Internat” from Assistance Publique - Hôpitaux de Paris. ALD is supported by the NIHR University College London Hospital (UCLH) Biomedical Research Centre (BRC). Work in AFP laboratory is supported by the European Research Council (ERC) under the European Union’s Horizon Europe research and innovation programme (grant agreement No. 101126209 “3D.FETOPRINT”). MP is supported by the NIHR Great Ormond Street Hospital Biomedical Research Centre (GOSH BRC). PDC is supported by the BREATH Consortium (552269), by the NIHR (NIHR-RP-2014-04-046), H2020 (668294, INTENS), OAK Foundation (W1095/OCAY-14-191), GOSH-CC (V5201) and the NIHR GOSH BRC. We thank Valerija Karaluka, Ayad Eddaoudi and the GOSICH facilities for the help with data acquisition. We are grateful to Ospedale Pediatrico Bambino Gesù, its Biobank service and the “cinque per mille” funds (Italian Ministry of Health) for their support. We thank Zeus Blacio Zambrano for helping with blinded images quantification. Finally, we are grateful to the patients and clinical teams at UCLH Fetal Medicine Unit, Professor Francesca Russo from Universitair Ziekenhuis Leuven and Professor Alexandra Benachi Hôpital Antoine Béclère – Assistance Publique – Hôpitaux de Paris and Dr Panicos Shangaris from Kings College Hospital who helped collecting the amniotic fluid samples used in this study.

## Author contribution

MFMG conceived and designed the study together with GDA, and with support and mentoring from PDC. GDA, GJZ and G Calà outlined and executed the experiments, analysed data and prepared the figures with help from GMC, AA, MM, RMdS and G Cenedese, under MFMG supervision. AM conducted the bioinformatics analysis. CC, MP and AFP helped conducting the pilot experiments and provided guidance and training to the team. ALD provided all the amniotic fluid samples utilised in the study, and managed the research midwives, the collection and consent procedures. SE helped with defining sample size and provided statistical support in the experimental design. MFMG, GDA, GJZ, G Calà and PDC drafted the manuscript. All the other authors provided contribution to the writing and approved to the final draft.

---

**Supplementary figure 1:**
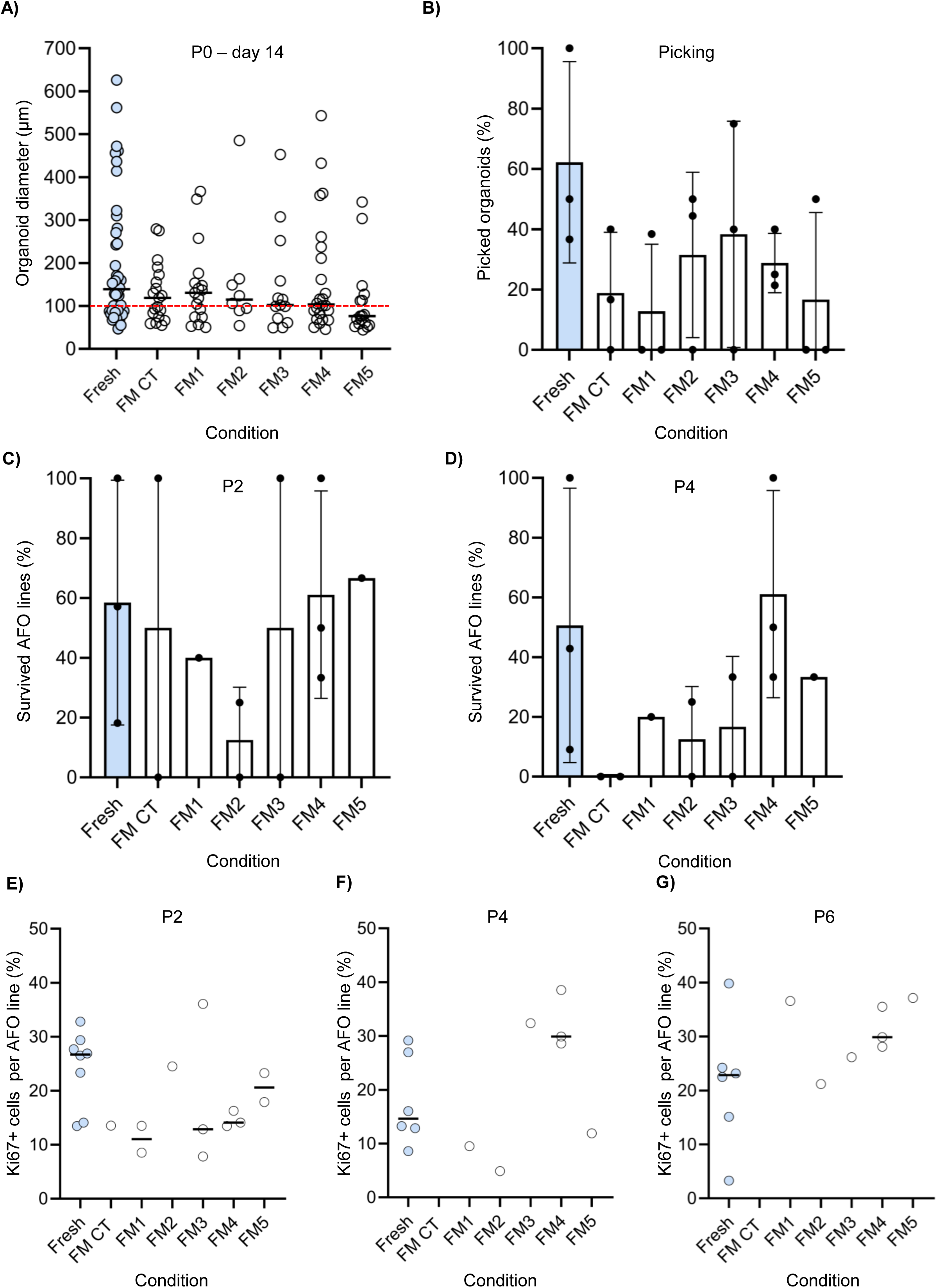
Characterisation of AFOs derived through Strategy 1. (A) Diameter of AFOs at passage 0, day 14 (each dot represents 1 organoid from n = 4 independent biological samples, mean is shown). Fresh vs FM comparison was performed with one-way ANOVA (non-significant); (B) Percentage of organoids picked at day 14 post-seeding. All organoids with diameter >= 100 µm were picked in n = 3 independent biological samples. mean ± s.e.m-; (C) and (D) percentage of AFO lines that could be expanded up to passage 2 and passage 4, relative to the number of picked organoids. mean ± s.e.m.;(E), (F) and (G) Quantification of Ki67 positive cells at passage 2, 4 and 6. Each dot represent 1 AFO line, mean is shown.

**Supplementary figure 2:**
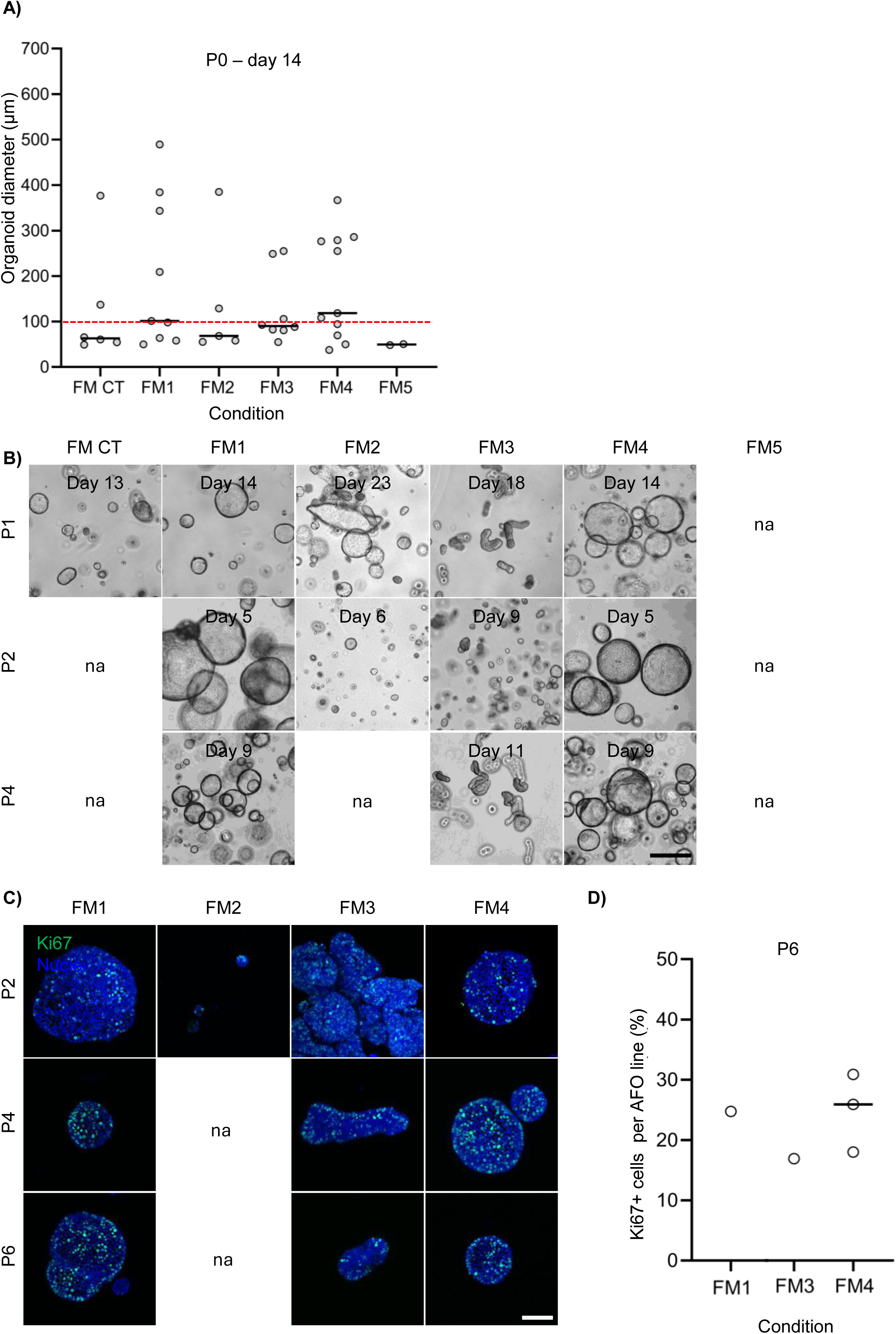
Characterisation of AFOs derived through Strategy 2. (A) Diameter of AFOs at passage 0, day 14. Longest diameter was used for asymmetric organoids. Measurement was performed using ImageJ software (each dot represents 1 organoid from n=4 independent biological samples, mean is shown). Fresh vs FM comparison was performed with one-way ANOVA (non-significant); (B) Brightfield images of expanding AFOs from thawed cells at passage 1, passage 2 and passage 4 (scale bar = 100 µm). (C) whole mount immunofluorescence staining of proliferation marker Ki67 counterstained with Hoechst (scale bar = 100 µm). (D) percentage of AFO lines that could be expanded up to passage 6, relative to the number of picked organoids (mean is shown).

## References

1. Corrò, C., Novellasdemunt, L. & Li, V. W. A brief history of organoids. Am J Physiol Cell Physiol 319, 151–165 (2020).

2. Zhao, Z. et al. Organoids. Nature Reviews Methods Primers 2, (2022).

3. Soto-Gamez, A., Gunawan, J. P., Barazzuol, L., Pringle, S. & Coppes, R. P. Organoid-based personalized medicine: from tumor outcome prediction to autologous transplantation. Stem Cells 42, 499 (2024).

4. Calà, G., Sina, B., De Coppi, P., Giobbe, G. G. & Gerli, M. F. M. Primary human organoids models: Current progress and key milestones. Frontiers in Bioengineering and Biotechnology vol. 11 Preprint at 10.3389/fbioe.2023.1058970 (2023).

5. Gerli, M. F. M. et al. Single-cell guided prenatal derivation of primary fetal epithelial organoids from human amniotic and tracheal fluids. Nat Med 30, 875– 887 (2024).

6. Atala, A. et al. The need for an organoid manufacturing, preservation, and distribution center. Stem Cells Transl Med 14, (2025).

7. Arutyunyan, I., Fatkhudinov, T. & Sukhikh, G. Umbilical cord tissue cryopreservation: A short review. Stem Cell Research and Therapy vol. 9 Preprint at 10.1186/s13287-018-0992-0 (2018).

8. Yoshizawa, R. S. Review: Public perspectives on the utilization of human placentas in scientific research and medicine. Placenta 34, 9–13 (2013).

9. Ballen, K. K., Gluckman, E. & Broxmeyer, H. E. Umbilical cord blood transplantation: the first 25 years and beyond. Blood 122, 491–498 (2013).

10. Broxmeyer, H. E. et al. Hematopoietic stem/progenitor cells, generation of induced pluripotent stem cells, and isolation of endothelial progenitors from 21-to 23.5-year cryopreserved cord blood. Blood 117, 4773–4777 (2011).

11. Castaneda, D. C. et al. Protocol for establishing primary human lung organoid-derived air-liquid interface cultures from cryopreserved human lung tissue. STAR Protoc 4, (2023).

12. Tsai, Y. H. et al. A Method for Cryogenic Preservation of Human Biopsy Specimens and Subsequent Organoid Culture. Cell Mol Gastroenterol Hepatol 6, 218 (2018).

13. Bui, B. N. et al. Organoids can be established reliably from cryopreserved biopsy catheter-derived endometrial tissue of infertile women. Reprod Biomed Online 41, 465–473 (2020).

14. Xue, W. et al. Effective cryopreservation of human brain tissue and neural organoids. Cell Reports Methods 4, 100777 (2024).

15. Heidari-Khoei, H. et al. Derivation of hormone-responsive human endometrial organoids and stromal cells from cryopreserved biopsies. Exp Cell Res 417, 113205 (2022).

16. Walsh, A. J., Cook, R. S., Sanders, M. E., Arteaga, C. L. & Skala, M. C. Drug response in organoids generated from frozen primary tumor tissues OPEN. (2015) doi:10.1038/srep18889.

17. Beato, F. et al. Establishing a Living Biobank of Patient-Derived Organoids of Intraductal Papillary Mucinous Neoplasms of the Pancreas. Lab Invest 101, 204 (2020).

18. Chen, P. et al. Establishing a cryopreserved biobank of living tumor tissues for drug sensitivity testing. Bioact Mater 46, 582–596 (2025).

19. Urbano, P. C. M., Angus, H. C. K., Gadeock, S., Schultz, M. & Kemp, R. A. Assessment of source material for human intestinal organoid culture for research and clinical use. BMC Res Notes 15, 1–8 (2022).

20. Calà Giuseppe et al. Derivation, expansion and cryopreservation of primary fetal organoids from second and third trimester human amniotic fluid cells. Nat Protoc.

21. de Coppi, P. et al. Regenerative medicine: prenatal approaches. The Lancet Child and Adolescent Health vol. 6 643–653 Preprint at 10.1016/S2352-4642(22)00192-4 (2022).

22. Verstegen, M. M. A. et al. Clinical applications of human organoids. Nat Med 31, 409–421 (2025).

